# RELAY: Relational In-Context Learning for Cellular Perturbation Prediction

**DOI:** 10.64898/2026.10.31.763027

**Authors:** Yuze Sun, Jinghao Feng, Yanfeng Wang, Weidi Xie, Ya Zhang, Xiaoman Zhang

**Author notes:** Corresponding author Weidi Xie; Ya Zhang, Xiaoman Zhang.

## Abstract

Cellular responses to perturbations vary across multiple experimental variables, such as perturbation identity, cell type, donor, and dose, defining a multidimensional condition space in which each measured population occupies one coordinate. Existing methods predict unmeasured conditions using explicit representations of these variable values, generalizing unseen values therefore depends on the availability and quality of such representations. We present **RELAY**, a relational in-context model that predicts unmeasured coordinates from measured responses at related coordinates. For each target, RELAY uses measured control–perturbed pairs from conditions that share one or more experimental variables with the target, such as the same perturbation in another cell type. The model receives only which variables are shared, not their values, and uses these demonstrations to predict the target response. We evaluate RELAY on chemical, genetic, and cytokine perturbation datasets under two settings: **unseen combinations**, where test values are individually observed during training but not in their target combinations, and **unseen variable values**, where test values are introduced only through context at inference. RELAY outperforms existing in-context methods on unseen variable values and supervised perturbation-prediction methods on unseen combinations. All resources will be open sourced^1 2^, including code and data.

## 1 Introduction

Accurately predicting cellular responses to perturbations at scale has the potential to reshape target discovery and therapeutic development (Bunne et al., 2024; Rood et al., 2024; Ji et al., 2021; Qi & Chapfuwa, 2026; Dibaeinia et al., 2026). These responses are jointly determined by multiple experimental variables: perturbation identity, such as a compound, cytokine, or genetic intervention; perturbation parameters, such as dose, strength, or duration; and biological context, including cell type, donor, and tissue. These factors define a multidimensional condition space in which each experimentally measured cell population occupies a single coordinate, and the space grows multiplicatively in the number of variables, so only a small fraction of it can ever be measured. Perturbation prediction is therefore fundamentally a problem of predicting responses at unmeasured coordinates from measurements elsewhere in this space.

Existing methods predict at a new coordinate by conditioning on the values of its variables. Perturbations may be represented by learned identifier embeddings, molecular structures, gene embeddings, or knowledge-graph features, while biological covariates are commonly encoded using categorical or learned representations (Lotfollahi et al., 2019; 2023; Roohani et al., 2024; Feng et al., 2026; Guo et al., 2026; Jiang et al., 2026). This paradigm works naturally when the relevant values and representations are available during training, but becomes more restrictive for unseen values.

Learned identifier embeddings cannot represent values absent from training, while descriptor-based approaches rely on external priors whose coverage and relevance vary across variable types. More-over, many experimental variables, such as donor identity or study-specific biological covariates, lack standardized descriptors (Kedzierska et al., 2025; Ahlmann-Eltze et al., 2025; Wu et al., 2026). Thus, generalization to unseen conditions often depends on how well each variable value can be represented.

Measured cell populations provide another way to specify a condition: through demonstrations of how cells actually respond, rather than through a predefined representation of the condition itself. Recent work shows that such in-context prediction is viable in single-cell biology. Stack (Dong et al., 2026) applies tabular attention over cell sets so that one population specifies a condition applied to another; scGen (Lotfollahi et al., 2019) transfers perturbation-induced shifts estimated from measured control and perturbed populations across cell types in a variational latent space; and general-purpose tabular foundation models with no biological pre-training are competitive with specialized architectures on several perturbation benchmarks (Palla et al., 2026; Hollmann et al., 2025; Qu et al., 2025; Zhu et al., 2025). These results suggest that measured responses can serve as informative conditioning signals for perturbation prediction. However, existing approaches largely treat context experiments as generic observations, without explicitly representing how each experiment is related to the target condition. This relation is informative because different context experiments reveal different components of the target coordinate. An experiment sharing the target perturbation shows how that perturbation acts in another biological context, whereas one sharing the target cell type shows how that cellular context responds to another perturbation.

In this paper, we make this relation explicit. For a target coordinate, **RELAY** receives measured control–perturbed pairs from conditions that share one or more variable values with the target, together with their shared-variable relations. Variable values are used only to identify matching demonstrations and are never encoded as model inputs. We implement this formulation with a shared pair en-coder for control–perturbed population pairs, relation embeddings that encode which variables each demonstration shares with the target, and target-to-context cross-attention that predicts expression changes in target control cells. The formulation extends naturally to multiple experimental variables by representing each demonstration according to the subset of variables it shares with the target.

We partition the condition space into base, context, and test regions (Figure 1), and evaluate two complementary settings. **Setting 1: unseen combinations**. Training uses both base and context conditions, so the values defining a test condition have been observed individually, but their combination has not. **Setting 2: unseen variable values**. Training uses only base conditions, so every value defining a test condition is unseen during training and is introduced only through context demonstrations at inference, with model parameters fixed. The first tests compositional generalization across previously observed values, whereas the second tests adaptation to entirely new variable values through inference-time evidence, with-out learning value-specific representations.

**Figure 1:**
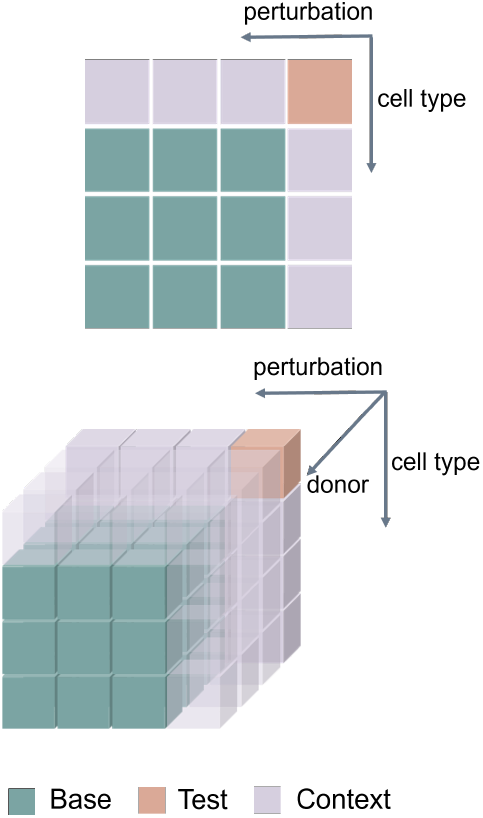
Partition of the condition space into base, context, and test conditions.

Our contributions are as follows.

- A relational formulation of perturbation prediction as in-context inference over a multidimensional condition space, using measured control–perturbed pairs that share variables with the target rather than explicit value representations.
- A unified in-context model that encodes control–perturbed pairs, represents their shared-variable relations to the target, and predicts target responses through cross-attention.
- An evaluation framework separating generalization to unseen combinations from generalization to unseen variable values, showing that RELAY improves perturbation-response prediction over supervised and in-context baselines under their respective settings.

## 2 Related Work

### Cellular perturbation prediction

Existing methods for cellular perturbation prediction span several modeling paradigms. Early approaches model perturbation effects in latent spaces or through compositional representations of perturbations and biological covariates (Lotfollahi et al., 2019; 2023). Other methods incorporate gene-level, molecular, or mechanistic priors to support generalization across perturbations (Roohani et al., 2024; Feng et al., 2026; Guo et al., 2026; Shi et al., 2026). More recent work models perturbation responses at the population level using optimal transport, flow matching, diffusion, or related distributional objectives (Bunne et al., 2023; Adduri et al., 2026; Klein et al., 2025; Yuan et al., 2026; He et al., 2026; Yu et al., 2026; Wang et al., 2026; Chen et al., 2026; Chi et al., 2026; Liu et al., 2026). Despite differences in architecture and training objective, these methods typically predict a target condition by conditioning on explicit representations of its perturbation and biological covariates.

### Context-based perturbation prediction

A complementary line of work uses measured cellular responses themselves as conditioning information. scGen transfers perturbation-induced latent shifts estimated from observed control and perturbed populations across cellular contexts (Lotfollahi et al., 2019). STACK (Dong et al., 2026) uses observed cell populations as in-context examples, while MapPFN (Sextro et al., 2026) and PerturbMap (Cui et al., 2026) leverage interventional observations or cross-context response transfer. General-purpose tabular foundation models likewise show that inference from context examples can be competitive on cellular perturbation benchmarks (Hollmann et al., 2025; Qu et al., 2025; Palla et al., 2026). These approaches demonstrate that measured responses can provide informative inference-time evidence. RELAY differs in explicitly representing how each demonstration is related to the prediction target: each control–perturbed population pair is encoded together with the subset of experimental variables it shares with the target, while the variable values themselves are never provided to the model.

## 3 Method

### 3.1 Problem Formulation

#### Problem Setup

We consider cellular perturbation experiments defined over a condition space with *K* variables, such as perturbation identity, dose, cell type, and donor. A condition is represented as *c* = (*c*_1_, … , *c*_*K*_) ∈ *C*, where *c*_*k*_ denotes the value of variable *k* and *C* is the space of all such conditions. Only a subset of conditions is experimentally measured. For a measured condition *c*, let 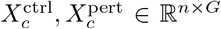,denote its control and perturbed cell populations over *G* genes, where *n* is the number of cells. Our goal is to predict 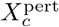 from its matched control population 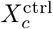 and measured responses at related conditions.

#### Experimental Settings

For each variable *k*, we partition its values into a base set and a held-out set. Among measured conditions, those composed entirely of base values form (*C*_base_), those composed entirely of held-out values form (*C*_test_), and those containing both base and held-out values form (*C*_ctx_) (Figure 1). We consider two settings:

- **Setting 1: unseen combinations**. Training uses *C* _base_ ∪*C* _ctx_: held-out variable values are individually observed during training, but their combinations are not.
- **Setting 2: unseen variable values**. Training uses only *C* _base_: every value composing a test condition is absent from training and is introduced only through context at inference.

#### Relational Context

Rather than encoding variable values, we specify the target through measured responses at related conditions. For a target condition *c* and a measured condition *c*′, define the set of shared variables as

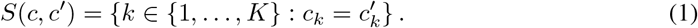

We use conditions satisfying ∅ ≠ *S*(*c, c*′) ⊊ {1, … , *K*} , so that each shares at least one variable value with the target while differing in at least one other variable. A *demonstration* consists of the measured control–perturbed pair at such a condition together with its relation to the target. For target *c*, we select *P* demonstration conditions and define *context* of the target as

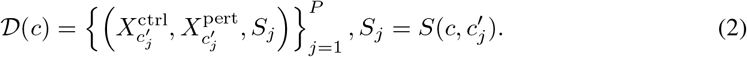

The prediction is

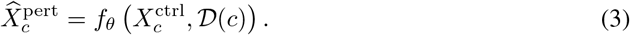

### 3.2 RELAY

Figure 2 provides an overview of **RELAY**. Given a target control population and a set of measured control–perturbed demonstrations, the model first encodes each demonstration with a shared pair encoder into a compact representation of its observed perturbation-induced transition (Section 3.2.1). It then incorporates the experimental variables shared between each demonstration and the target to form a relation-aware context memory (Section 3.2.2). Target control cells attend to this memory to integrate information across related conditions and predict perturbation-induced expression changes (Section 3.2.3). All components are trained jointly with population-level supervision, without requiring cell-to-cell correspondence between control and perturbed populations (Section 3.3).

**Figure 2:**
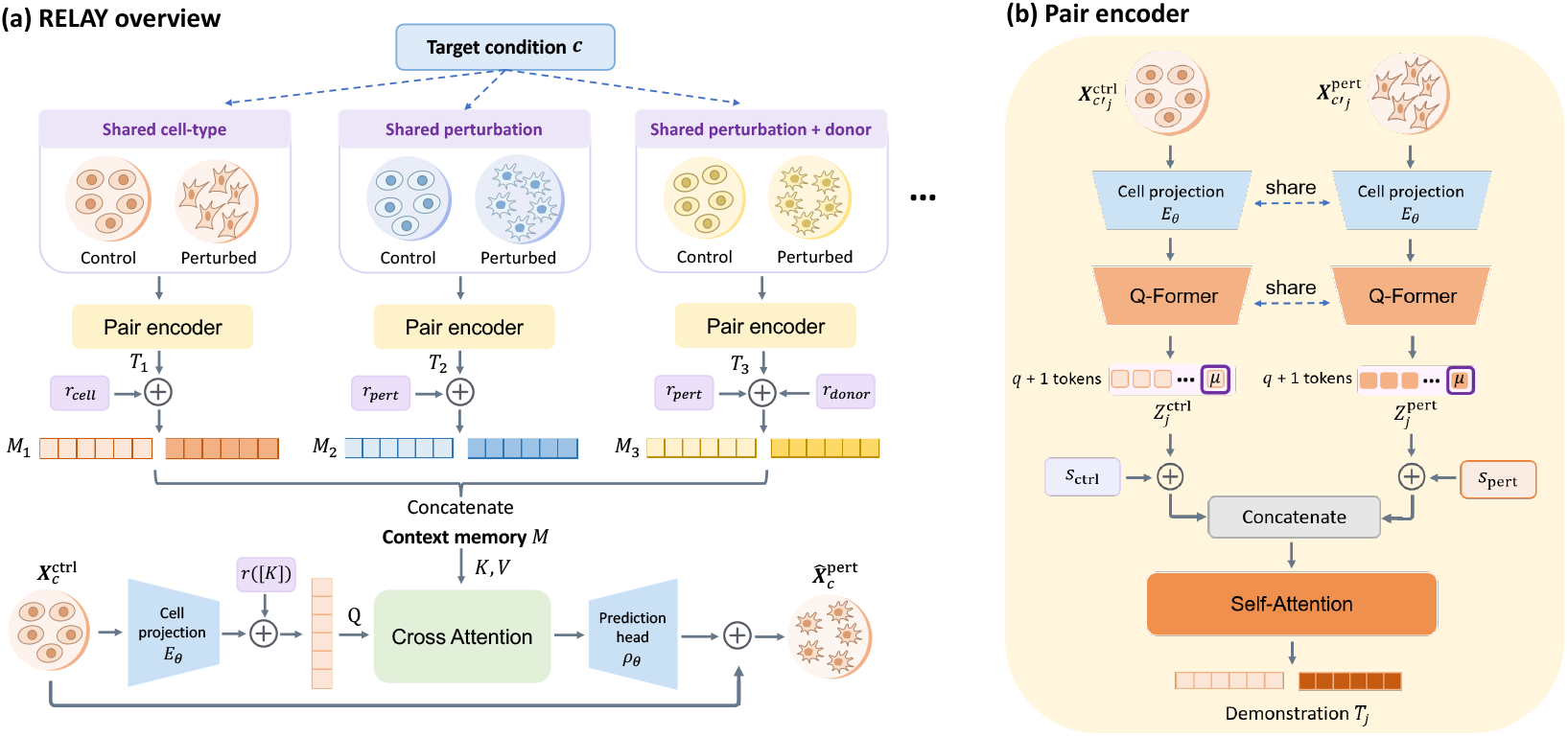
Overview of RELAY. **(a)** Target control cells attend to measured control–perturbed demonstrations to predict the target perturbation response, using shared-variable relations rather than explicit variable-value representations. **(b)** A shared pair encoder summarizes each control and perturbed population and jointly models the pair with self-attention, with state embeddings distinguishing the two population states.

#### 3.2.1. Demonstration Encoding

Each demonstration contains a control–perturbed cell population pair, which we encode with a pair encoder (Figure 2b). To represent multiple demonstrations efficiently, the pair encoder first compresses each cell population into a fixed number of tokens and then jointly encodes the two populations to capture the observed perturbation-induced transition. Specifically, each cell is mapped to a *d*-dimensional representation by a projection layer *E*_*θ*_ : ℝ^*G*^ →ℝ^*d*^. A Q-Former (Li et al., 2023) with *q* learned queries then summarizes each cell population into *q* tokens, to which we append a projected population-mean token. For demonstration *j*, this yields

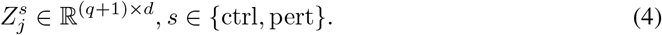

The cell projection and Q-Former are shared between the control and perturbed populations, and the resulting summaries are invariant to cell ordering.

To distinguish the control and perturbed populations, we add learnable state embeddings **s**_ctrl_, **s**_pert_ ∈ ℝ^*d*^ to their corresponding tokens. The two sets of tokens are then concatenated and jointly processed by *L*_*p*_ self-attention blocks:

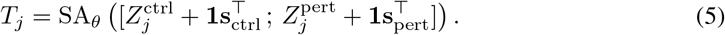

Here, [·;·] denotes concatenation along the token dimension, and **1** broadcasts the corresponding state embedding to all tokens. By jointly modeling the control and perturbed populations within each demonstration, the pair encoder output *T*_*j*_ represents its observed perturbation-induced transition. The same pair encoder is shared across all demonstrations.

#### 3.2.2. Relation Embedding

The demonstration representation *T*_*j*_ captures the observed control–perturbed transition, but does not specify how the demonstration is related to the target condition. We therefore encode which experimental variables are shared between each demonstration and the target. For demonstration *j*, let *S*_*j*_ denote the set of shared variables. Each variable *k* is associated with a learnable embedding *r*_*k*_ ∈ ℝ^*d*^, and the target–demonstration relation is represented as:

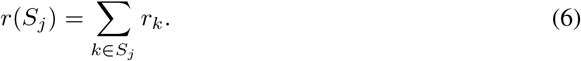

For example, if a demonstration shares perturbation identity and donor with the target, its relation embedding is *r*_pert_ + *r*_donor_.

We add this relation embedding to all tokens of the corresponding demonstration representation:

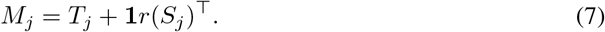

The representations of all the P demonstrations are then concatenated along the token dimension to form the context memory,

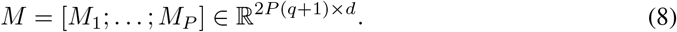

#### 3.2.3. Context-Conditioned Prediction

Given the context memory *M* , we use the target control cells as queries to attend to relevant information across demonstrations and predict the target perturbed population. Unlike the demonstration populations, which are compressed into a fixed number of tokens, we retain one query token for each target control cell so that the model directly produces a predicted expression profile for each cell. Specifically, the same projection layer *E*_*θ*_ maps the target control cells into *d*-dimensional representations. Since the target condition matches itself on all *K* variables, we add the corresponding relation embedding *r*([*K*]), yielding

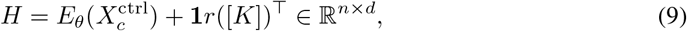

*H* = *E*_*θ*_(*X*^ctrl^) + **1***r*([*K*])^⊤^ ∈ ℝ^*n*×*d*^, (9) where [*K*] = {1, … , *K}* . This embedding distinguishes the target queries from demonstration tokens and represents the target as the reference condition.

The target queries are then updated through *L*_*t*_ cross-attention blocks over the context memory *M* , allowing each target cell to integrate information from the encoded demonstration transitions and their relations to the target. A prediction head *ρ*_*θ*_ : ℝ^*d*^ → ℝ^*G*^ maps the resulting representations to perturbation-induced expression changes. We predict the perturbed expression as a residual from the target control state:

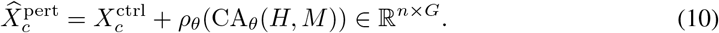

CA_*θ*_ denotes the stack of cross-attention blocks, including cross-attention, feed-forward layers, normalization, and residual connections. The target controls provide the baseline cell states, while the context memory informs how these states are expected to change under the target condition.

### 3.3. Training and Inference

We train all components jointly using an energy-kernel maximum mean discrepancy loss ℒ_MMD_:

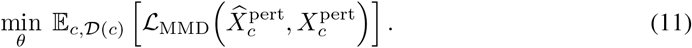

The population-level objective compares the predicted and observed distributions without requiring cell-to-cell correspondence. Architecture, hyperparameters, and optimization details are provided in Appendix B.2.

## 4 Experiments and Results

### 4.1 Experimental Setup

#### Benchmarks

We evaluate four benchmarks spanning chemical, genetic, and cytokine perturbations. **Tahoe-100M** (Zhang et al., 2025) varies compound and cell line, using 5 *µ*M dose and matched DMSO controls. **Replogle–Nadig** (Replogle et al., 2022; Nadig et al., 2024) varies CRISPRi target and cell line, with non-targeting guides as controls. From the Parse cytokine dataset (Oesinghaus et al., 2025), **PBMC-2D** varies cytokine and cell type within Donor9, while **PBMC-3D** additionally varies donor; both use matched PBS controls. Dataset sizes and condition-level cell counts are summarized in Appendix A.1.

#### Evaluation protocol

We follow the partitions and settings defined in Section 3.1. **Setting 1** trains on base and context conditions to predict unseen combinations, whereas **Setting 2** trains only on base conditions and introduces context at inference without parameter updates. Both settings use the same test conditions and inference context. On Tahoe-100M, base and held-out compounds are additionally separated by Bemis–Murcko scaffold (Bemis & Murcko, 1996). All methods use identical data splits and preprocessing, including library-size normalization to 10^4^, log transformation, and a common highly variable gene panel selected using training-visible cells only. Split construction and preprocessing are detailed in Appendices A and B.1, respectively.

#### Baselines

We evaluate broadly applicable baselines across chemical, genetic, and cytokine perturbation benchmarks. Setting 1 includes five supervised methods: **STATE** (Adduri et al., 2026) **models perturbation responses at the population level. CellFlow** (Klein et al., 2025) **learns covariate-conditioned transport maps from control to perturbed populations. CPA** (Lotfollahi et al., 2023) **combines a basal cell state with perturbation and covariate embeddings. We also include two recent baselines: scDFM** (Yu et al., 2026) generates perturbed cell distributions from control states and perturbation embeddings via conditional flow matching with an MMD objective. **PerturbDiff** (Yuan et al., 2026) models perturbation responses through functional diffusion over cell-population distributions embedded in a Hilbert space.

Setting 2 includes three context-based methods: **STACK** (Dong et al., 2026) uses attention over measured cell populations to predict responses in a target population. **TabPFN** (Hollmann et al., 2025; **Palla et al**., **2026)** uses a pretrained tabular model to infer target responses from context examples. **scGen** (Lotfollahi et al., 2019) is adapted to transfer perturbation-induced latent shifts from context populations to target controls. Both settings also include two non-learning baselines: **Untreated** directly uses target controls as the prediction. **Mean perturbation** adds the mean perturbed-minus-control shift for the same perturbation in other cellular contexts to target controls.

#### Metrics

We evaluate expression accuracy and perturbation-response recovery, averaging each metric equally over test conditions. **MSE** and **MAE** measure differences between predicted and observed mean expression profiles. **PDCorr** measures Pearson correlation coefficient between predicted and ground-truth relative perturbation effects. **PDS** measures perturbation specificity by ranking the correct measured response against other perturbations sharing the same non-perturbation variables. **REACH** is based on differential expression (DE) analysis and measures the fraction of reference DE genes recovered with reliable response directions, prioritizing genes with stronger predicted statistical evidence (Welch, 1947; Benjamini & Hochberg, 1995). Lower MSE and MAE and higher PDCorr, PDS, and REACH indicate better performance. Formal definitions of the training objective and evaluation metrics are provided in Appendix B.3.

### 4.2 Main Results

Tables 1 and 2 summarize performance in the two evaluation settings. We report means and standard deviations over three seeds.

**Table 1:**
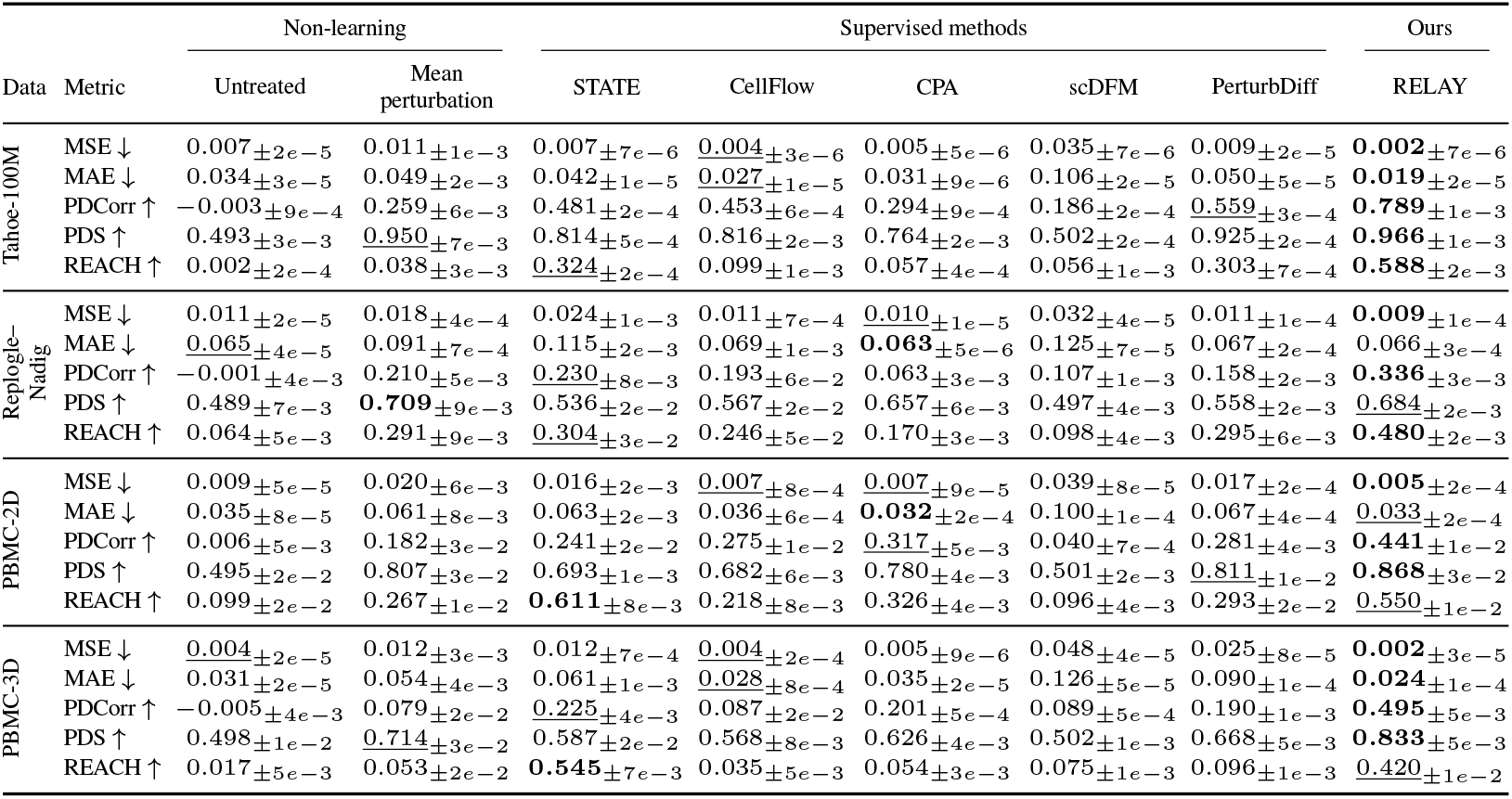
Comparison of methods for unseen combination prediction (Setting 1).

**Table 2:**
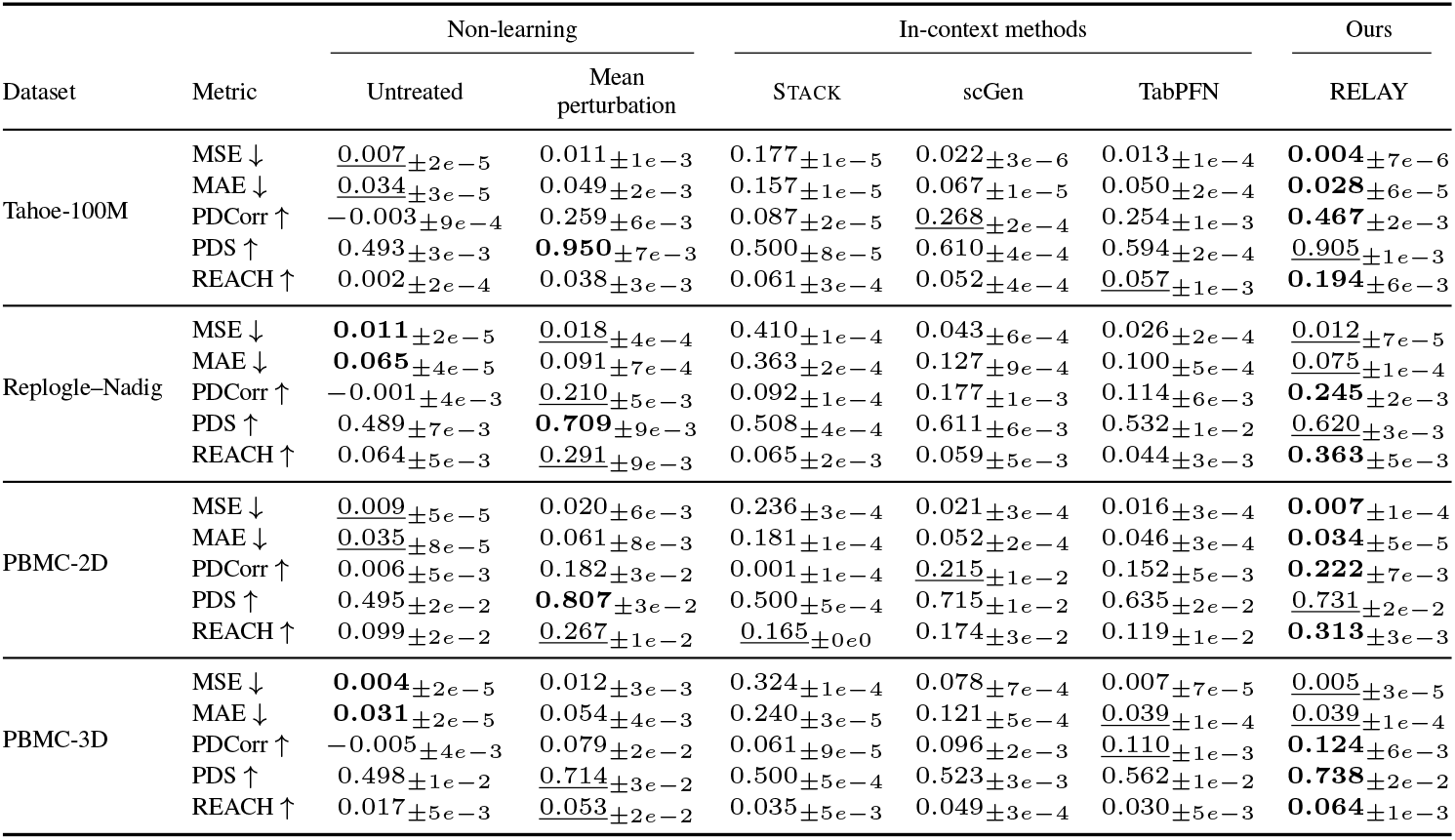
Comparison of methods for prediction with unseen variable values (Setting 2).

| Dataset | Metric | Non-learning |  | In-context methods |  |  | Ours |
|---|---|---|---|---|---|---|---|
|  |  | Untreated | Mean perturbation | Stack | scGen | TabPFN | RELAY |
| Tahoe-100M | MSE ↓ | <u>0.007</u><sub>±2e−5</sub> | 0.011<sub>±1e−3</sub> | 0.177<sub>±1e−5</sub> | 0.022<sub>±3e−6</sub> | 0.013<sub>±1e−4</sub> | **0.004**<sub>±7e−6</sub> |
|  | MAE ↓ | <u>0.034</u><sub>±3e−5</sub> | 0.049<sub>±2e−3</sub> | 0.157<sub>±1e−5</sub> | 0.067<sub>±1e−5</sub> | 0.050<sub>±2e−4</sub> | **0.028**<sub>±6e−5</sub> |
|  | PDCorr ↑ | −0.003<sub>±9e−4</sub> | 0.259<sub>±6e−3</sub> | 0.087<sub>±2e−5</sub> | <u>0.268</u><sub>±2e−4</sub> | 0.254<sub>±1e−3</sub> | **0.467**<sub>±2e−3</sub> |
|  | PDS ↑ | 0.493<sub>±3e−3</sub> | **0.950**<sub>±7e−3</sub> | 0.500<sub>±8e−5</sub> | 0.610<sub>±4e−4</sub> | 0.594<sub>±2e−4</sub> | <u>0.905</u><sub>±1e−3</sub> |
|  | REACH ↑ | 0.002<sub>±2e−4</sub> | 0.038<sub>±3e−3</sub> | 0.061<sub>±3e−4</sub> | 0.052<sub>±4e−4</sub> | <u>0.057</u><sub>±1e−3</sub> | **0.194**<sub>±6e−3</sub> |
| Replogle–Nadig | MSE ↓ | **0.011**<sub>±2e−5</sub> | <u>0.018</u><sub>±4e−4</sub> | 0.410<sub>±1e−4</sub> | 0.043<sub>±6e−4</sub> | 0.026<sub>±2e−4</sub> | <u>0.012</u><sub>±7e−5</sub> |
|  | MAE ↓ | **0.065**<sub>±4e−5</sub> | 0.091<sub>±7e−4</sub> | 0.363<sub>±2e−4</sub> | 0.127<sub>±9e−4</sub> | 0.100<sub>±5e−4</sub> | <u>0.075</u><sub>±1e−4</sub> |
|  | PDCorr ↑ | −0.001<sub>±4e−3</sub> | <u>0.210</u><sub>±5e−3</sub> | 0.092<sub>±1e−4</sub> | 0.177<sub>±1e−3</sub> | 0.114<sub>±6e−3</sub> | **0.245**<sub>±2e−3</sub> |
|  | PDS ↑ | 0.489<sub>±7e−3</sub> | **0.709**<sub>±9e−3</sub> | 0.508<sub>±4e−4</sub> | 0.611<sub>±6e−3</sub> | 0.532<sub>±1e−2</sub> | <u>0.620</u><sub>±3e−3</sub> |
|  | REACH ↑ | 0.064<sub>±5e−3</sub> | <u>0.291</u><sub>±9e−3</sub> | 0.065<sub>±2e−3</sub> | 0.059<sub>±5e−3</sub> | 0.044<sub>±3e−3</sub> | **0.363**<sub>±5e−3</sub> |
| PBMC-2D | MSE ↓ | <u>0.009</u><sub>±5e−5</sub> | 0.020<sub>±6e−3</sub> | 0.236<sub>±3e−4</sub> | 0.021<sub>±3e−4</sub> | 0.016<sub>±3e−4</sub> | **0.007**<sub>±1e−4</sub> |
|  | MAE ↓ | <u>0.035</u><sub>±8e−5</sub> | 0.061<sub>±8e−3</sub> | 0.181<sub>±1e−4</sub> | 0.052<sub>±2e−4</sub> | 0.046<sub>±3e−4</sub> | **0.034**<sub>±5e−5</sub> |
|  | PDCorr ↑ | 0.006<sub>±5e−3</sub> | 0.182<sub>±3e−2</sub> | 0.001<sub>±1e−4</sub> | <u>0.215</u><sub>±1e−2</sub> | 0.152<sub>±5e−3</sub> | **0.222**<sub>±7e−3</sub> |
|  | PDS ↑ | 0.495<sub>±2e−2</sub> | **0.807**<sub>±3e−2</sub> | 0.500<sub>±5e−4</sub> | 0.715<sub>±1e−2</sub> | 0.635<sub>±2e−2</sub> | <u>0.731</u><sub>±2e−2</sub> |
|  | REACH ↑ | 0.099<sub>±2e−2</sub> | <u>0.267</u><sub>±1e−2</sub> | <u>0.165</u><sub>±0e0</sub> | 0.174<sub>±3e−2</sub> | 0.119<sub>±1e−2</sub> | **0.313**<sub>±3e−3</sub> |
| PBMC-3D | MSE ↓ | **0.004**<sub>±2e−5</sub> | 0.012<sub>±3e−3</sub> | 0.324<sub>±1e−4</sub> | 0.078<sub>±7e−4</sub> | 0.007<sub>±7e−5</sub> | <u>0.005</u><sub>±3e−5</sub> |
|  | MAE ↓ | **0.031**<sub>±2e−5</sub> | 0.054<sub>±4e−3</sub> | 0.240<sub>±3e−5</sub> | 0.121<sub>±5e−4</sub> | <u>0.039</u><sub>±1e−4</sub> | <u>0.039</u><sub>±1e−4</sub> |
|  | PDCorr ↑ | −0.005<sub>±4e−3</sub> | 0.079<sub>±2e−2</sub> | 0.061<sub>±9e−5</sub> | 0.096<sub>±2e−3</sub> | <u>0.110</u><sub>±1e−3</sub> | **0.124**<sub>±6e−3</sub> |
|  | PDS ↑ | 0.498<sub>±1e−2</sub> | <u>0.714</u><sub>±3e−2</sub> | 0.500<sub>±5e−4</sub> | 0.523<sub>±3e−3</sub> | 0.562<sub>±1e−2</sub> | **0.738**<sub>±2e−2</sub> |
|  | REACH ↑ | 0.017<sub>±5e−3</sub> | <u>0.053</u><sub>±2e−2</sub> | 0.035<sub>±5e−3</sub> | 0.049<sub>±3e−4</sub> | 0.030<sub>±5e−3</sub> | **0.064**<sub>±1e−3</sub> |

### Setting 1: unseen combinations

RELAY achieves the highest PDCorr and lowest MSE on all four benchmarks, while also outperforming all supervised baselines in PDS. The largest absolute PDCorr margin over the strongest supervised baseline occurs on PBMC-3D, where RELAY reaches 0.495 compared with 0.225 for STATE, highlighting its effectiveness in a condition space spanning perturbation, cell type, and donor. Beyond these consistent gains, RELAY also achieves the highest REACH on Tahoe-100M and Replogle–Nadig, supporting its ability to recover differential-expression response directions. These results show that relational context supports generalization to unobserved combinations of previously seen variable values, improving both perturbation-effect recovery and mean expression accuracy.

### Setting 2: unseen variable values

RELAY outperforms the learned context-based baselines STACK, scGen, and TabPFN in PDCorr, PDS, and REACH on all four benchmarks. The largest PDCorr margin occurs on Tahoe-100M, where RELAY reaches 0.467 compared with 0.268 for sc-Gen. Performance is generally lower than in Setting 1 when the target variable values are entirely absent from training, reflecting the greater difficulty of inference from context alone. Simple base-lines nevertheless remain competitive on some metrics: Mean perturbation achieves the highest PDS on the three 2D benchmarks, while Untreated obtains lower MSE and MAE than RELAY on Replogle–Nadig and PBMC-3D despite near-zero PDCorr. This contrast highlights that low absolute expression error does not necessarily imply accurate recovery of perturbation effects.

### 4.3 Ablation and Model Analysis

We conduct the following analyses on PBMC-3D under Setting 1, except where noted otherwise.

#### State and relation embeddings

Combining both embeddings gives the highest mean scores on all three metrics (Figure 3a): PDCorr, PDS, and REACH reach 0.495, 0.833, and 0.420, respectively. Removing relation embedding reduces these values to 0.170, 0.599, and 0.040, while removing state embedding gives 0.303, 0.582, and 0.162. Removing both reduces performance further to 0.096, 0.507, and 0.026. These results suggest that state and relation embeddings capture complementary aspects of the context, with relation embeddings contributing more strongly to PDCorr and REACH.

**Figure 3:**
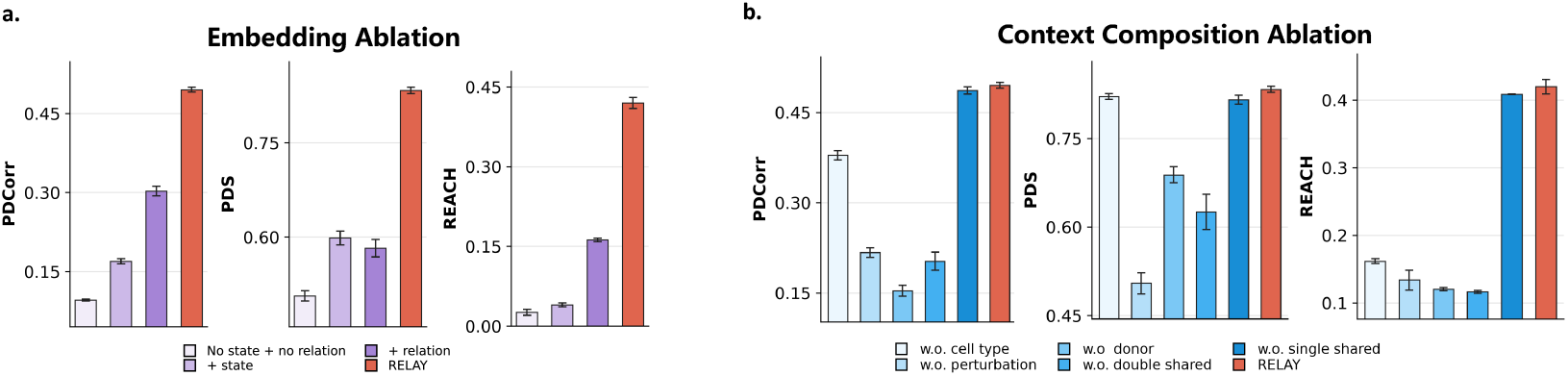
Effects of model components and context composition. (a) Removal of state and relation embeddings. (b) Removal of context groups defined by the variables shared with the target. Bars show means; error bars indicate standard deviations.

#### Context composition

Context groups contribute differently across metrics (Figure 3b). Removing perturbation-matching groups brings PDS close to chance (0.505), whereas removing donor-matching groups causes the largest PDCorr drop (to 0.154). Removing double-shared groups is more damaging than removing single-shared groups: retaining only single-shared groups gives PDCorr 0.203, PDS 0.626, and REACH 0.116, whereas retaining only double-shared groups gives 0.486, 0.816, and 0.409. These results show that both the identity and the number of target-matching variables determine context utility, while the retained performance with missing groups suggests applicability when related measurements are sparse.

#### Context size

Here, *m*_1_ and *m*_2_ denote the numbers of demonstrations sampled per context group sharing exactly one and two target variables, respectively. The results (Figure 4a)show that performance is driven primarily by *m*_2_, whereas changing *m*_1_ produces only small differences. At fixed *m*_1_, increasing *m*_2_ from 1 to 5 raises PDCorr, PDS, and REACH by roughly 0.075, 0.053, and 0.107, respectively; increasing *m*_2_ from 5 to 9 adds only about 0.012, 0.010, and 0.022. Under comparable context budgets, allocating more examples to double-shared groups is therefore preferable, although the gains become small once *m*_2_ ≥ 5.

**Figure 4:**
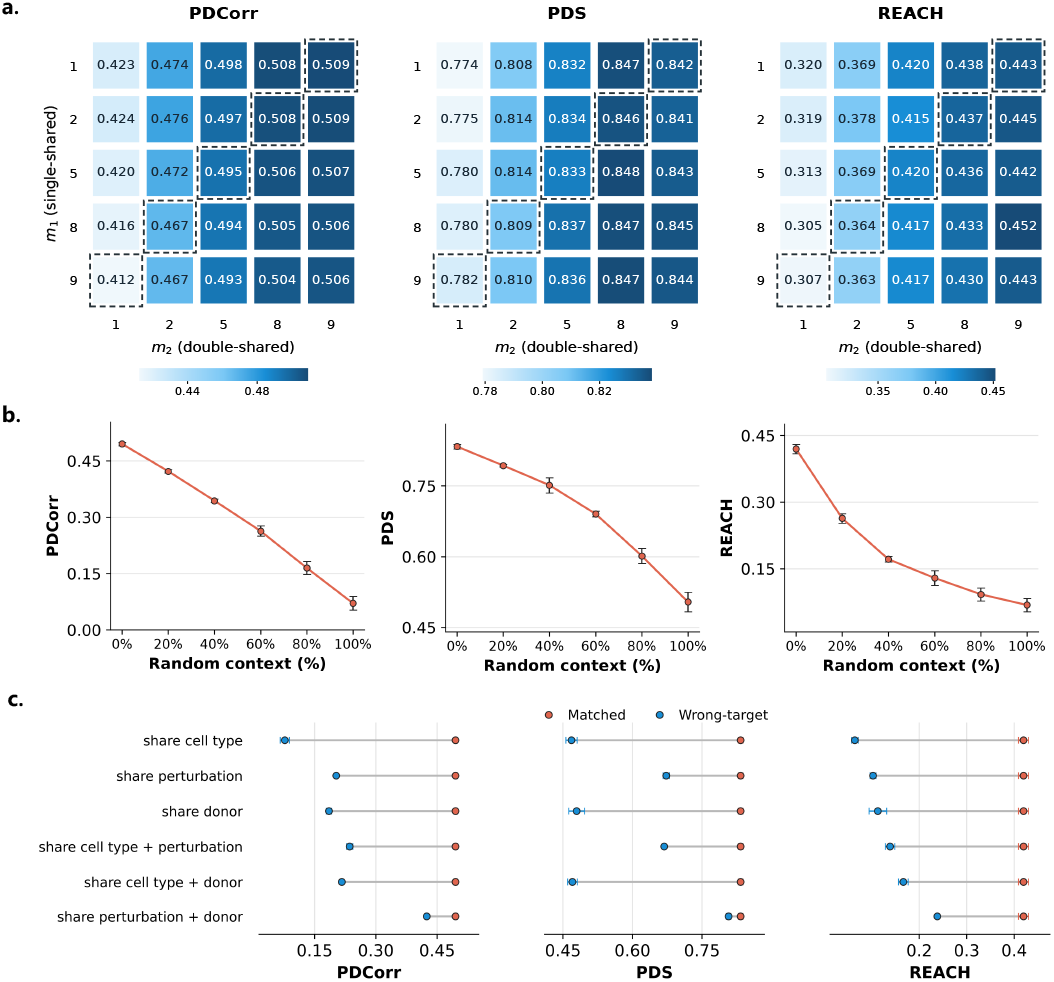
Sensitivity to context quantity and relevance. (a) Varying the numbers of demonstrations per single-shared group (*m*_1_) and double-shared group (*m*_2_). (b) Random replacement of context pairs. (c) Replacement with the complete context of another target sharing the listed variables.

#### Random context replacement

To test whether the model relies on the correspondence between the provided context and the target condition, we replace demonstrations at inference while keeping the target controls, context size, and group sizes unchanged (Figure 4b). Random replacement preserves each control–perturbed pair but leaves the original relation embeddings unchanged, disrupting the correspondence between demonstrations and the target. All three metrics decrease monotonically with the replacement fraction: PDCorr falls from 0.495 to 0.071, PDS from 0.833 to 0.504, and REACH from 0.420 to 0.048. These results confirm that the model is sensitive to the correspondence between demonstrations and the target condition.

#### Borrowed context replacement

Borrowing the complete context of another target also lowers all three metrics for every tested overlap pattern. Sharing both perturbation and donor best preserves all three metrics among these replacements, reaching PDCorr 0.424, PDS 0.807, and REACH 0.238, but still does not recover matched-context performance (Figure 4c). Together with random replacement, these results show that RELAY retains useful predictive signal under partially noisy or mismatched context, while accurate target-specific demonstration selection remains important.

#### Attention to context

We examine which context populations receive more attention from target control cells, with each context group containing 5 demonstrations (Figure 5). In PBMC-2D, attention favors control-origin tokens in demonstrations sharing the target cell type, but perturbed-origin tokens in those sharing the target perturbation. PBMC-3D shows a similar pattern: demonstrations sharing both cell type and donor receive more attention on control-origin tokens, whereas those sharing perturbation alone or perturbation and donor receive more attention on perturbed-origin tokens. Thus, the relative attention to the two populations within a pair varies with the variables shared between the demonstration and the target. In PBMC-3D, two-variable matches also receive more attention than single-variable matches despite equal group sizes, consistent with the larger performance drops observed when two-variable matches are removed. These observations describe how attention is distributed, rather than establish which demonstrations cause better predictions. More-over, control- and perturbed-origin tokens are not independent representations of the two states, as pair self-attention allows them to exchange information.

**Figure 5:**
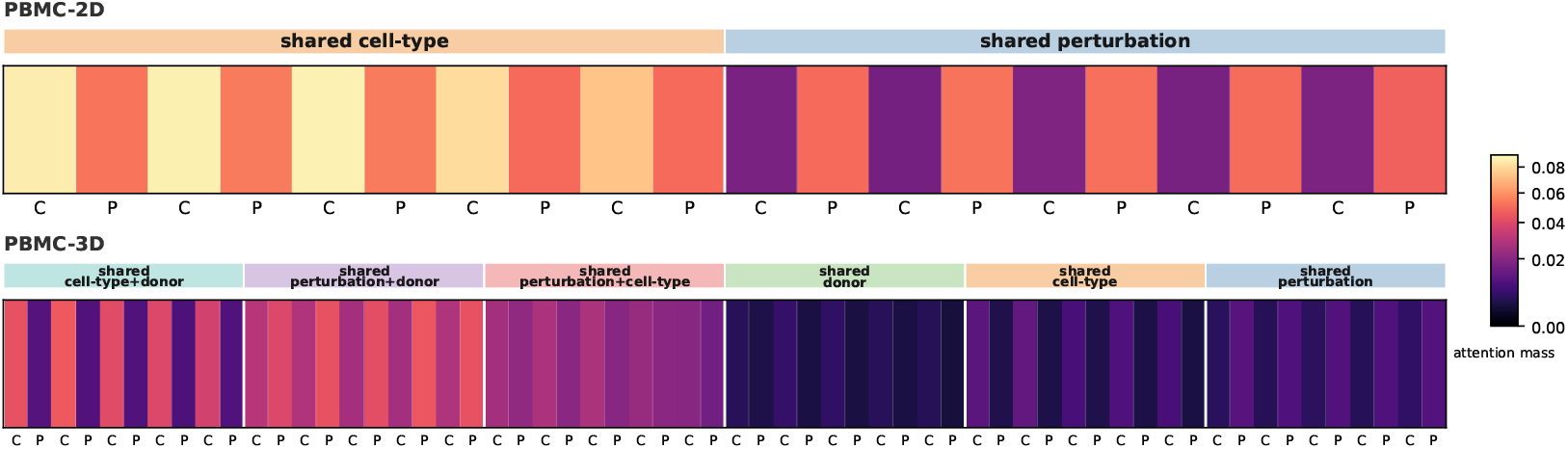
Cross-attention to context in PBMC-2D (top) and PBMC-3D (bottom), averaged over 256 samples. C/P denote control/perturbed tokens; group labels indicate target-shared variables. Colors show attention mass summed over each population’s compressed tokens.

## 5 Discussion and Conclusion

RELAY predicts cellular perturbation responses from measured control–perturbed demonstrations, using their shared-variable relations to the target rather than explicit representations of variable values. This formulation supports prediction of unseen combinations and adaptation to unseen values through inference-time context. Pair self-attention and the energy-distance MMD both operate on cell populations, and the loss compares full cell sets rather than means, discouraging collapse to an averaged response. Broader evaluation is also constrained by the scarcity of high-quality perturbation datasets with sufficient cells per condition and dense coverage of multidimensional spaces that include dose and time. Such datasets would enable stronger tests of relational generalization beyond the variables studied here and support extensions from exact-match relations to graded relations over continuous experimental variables.

### Reproducibility Statement

The paper specifies the model architecture, training objective, data preprocessing, evaluation protocol, and the two generalization settings. The public source datasets are identified in the Experimental Setup (Zhang et al., 2025; Replogle et al., 2022; Nadig et al., 2024; Oesinghaus et al., 2025). The appendix describes the condition-space partitions, base/context/test subsets, entropy-gap statistics, normalization and HVG selection, principal hyperparameters, ablations, and evaluation metrics. All reported aggregate results use the stated split definitions, and the same preprocessing and metric conventions across methods. The final release will provide the source code, fixed data split manifests, model checkpoints, training configurations, and evaluation scripts in a public repository.

## Ethics Statement

This study relies solely on publicly accessible single-cell gene expression datasets, with all data used in compliance with the applicable data use agreements. No directly identifiable personal information was used in the analyses. The methods presented here are designed to support computational modeling and hypothesis generation in biological research. Although these methods may eventually contribute to drug discovery or therapeutic design, they are not intended for direct clinical application. We advocate the responsible and transparent use of generative models in biological research.

### AI Use Statement

In this work, we used generative AI tools to assist with software implementation, literature search and summarization, figure preparation, and manuscript editing. They were also used to provide feedback on experimental design, refine hypotheses, and assist with the interpretation and presentation of results. The core research questions, methodological ideas, model design, experiments, and scientific conclusions were developed by the authors. All AI-assisted outputs were reviewed and verified by the authors, who take full responsibility for the final content of this work.

## A Split Construction and Entropy Gaps

### A.1 Dataset Scale

We summarize the benchmark subsets used in our experiments. Table 3 reports the number of base and held-out values for each variable. Table 4 reports the number of measured conditions and the median number of treated cells per condition in *C*_base_, *C*_ctx_, and *C*_test_.

**Table 3:** Numbers of base / held-out variable values. Cell identity denotes cell line for Replogle– Nadig and Tahoe-100M, and cell type for PBMC. PBMC-2D fixes Donor9; Tahoe-100M fixes drug dose at 5 *µ*M.

| Dataset | Perturbation | Cell identity | Donor |
|---|---|---|---|
| Replogle–Nadig | 1,642 / 411 | 3 / 1 | None |
| Tahoe-100M | 303 / 76 | 40 / 10 | None |
| PBMC-2D | 72 / 18 | 14 / 4 | Fixed (Donor9) |
| PBMC-3D | 72 / 18 | 14 / 4 | 10 / 2 |

**Table 4:**
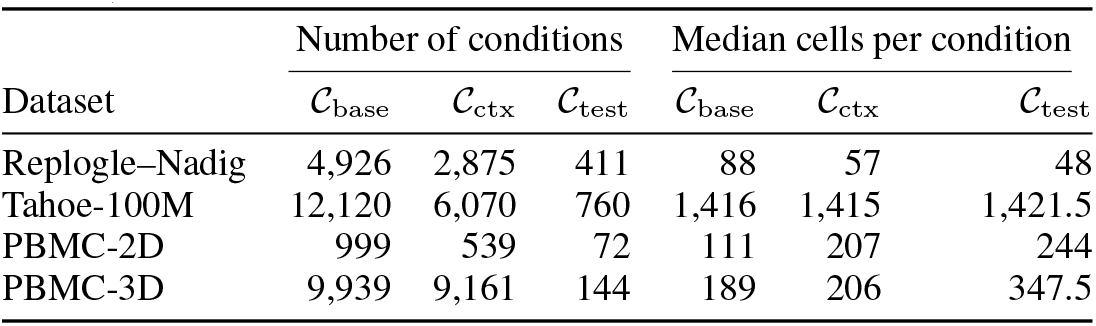
Measured condition counts and median treated-cell counts per condition. Only nonempty conditions are included.

|  | Number of conditions |  |  | Median cells per condition |  |  |
|---|---|---|---|---|---|---|
| Dataset | 𝒞<sub>base</sub> | 𝒞<sub>ctx</sub> | 𝒞<sub>test</sub> | 𝒞<sub>base</sub> | 𝒞<sub>ctx</sub> | 𝒞<sub>test</sub> |
| Replogle–Nadig | 4,926 | 2,875 | 411 | 88 | 57 | 48 |
| Tahoe-100M | 12,120 | 6,070 | 760 | 1,416 | 1,415 | 1,421.5 |
| PBMC-2D | 999 | 539 | 72 | 111 | 207 | 244 |
| PBMC-3D | 9,939 | 9,161 | 144 | 189 | 206 | 347.5 |

### A.2 Split Construction and Balance

#### Binning strategy

We rank variable values by their treated-cell counts and divide them into bins containing nearly equal numbers of values, then sample held-out values within each bin. All four datasets use six perturbation bins; Tahoe-100M and both PBMC datasets additionally use four cell-line or cell-type bins, and PBMC-3D uses two donor bins. Replogle–Nadig holds out approximately 20% of perturbations and one cell line (HepG2). Tahoe-100M targets a 20% holdout on both axes while keeping drug scaffold groups intact, with a penalty for deviations from the target bin allocation. Both PBMC splits hold out three perturbations per bin and one cell type per bin; PBMC-3D also holds out one donor per bin. PBMC-2D searches donor-specific candidates and selects Donor9. Bins stratify sampling by abundance; entropy is computed over individual variable values or joint conditions, not over the bins.

#### Entropy gaps

For a positive count vector **n** = (*n*_1_, … , *n*_*K*_), define *p*_*i*_ = *n*_*i*_*/ ∑*_*j*_ *n*_*j*_ and the normalized entropy imbalance

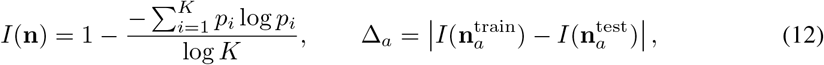

Zero counts are excluded and *I* = 0 when *K*≤ 1, including the single held-out cell line in Replogle– Nadig.*A* Here contains each variable axis and the joint pair (2D) or triplet (3D) distribution. For Replogle–Nadig and PBMC, marginal counts are totaled over the full treated subset and restricted to train/test variable values; Tahoe uses marginals within the respective train/test target blocks. Joint counts always come from those target blocks. Each table entry is one such absolute train–test entropy difference; smaller values indicate more similar abundance imbalance. The gaps are used as one component of split selection together with count-distribution and support criteria, rather than as a model-training loss. Tables 5 and 6 report the selected splits.

**Table 5:** Entropy gaps for the three 2D benchmarks. Cell identity denotes cell line for Replogle–Nadig and Tahoe-100M, and cell type for PBMC. Joint denotes perturbation × cell identity. Values are fixed-split statistics, rounded to six decimal places; smaller gaps indicate more similar normalized entropy imbalance.

| Dataset | Perturbation | Cell identity | Joint |
|---|---|---|---|
| Replogle–Nadig | 0.007273 | 0.015557 | 0.009409 |
| Tahoe-100M | 0.007415 | 0.007561 | 0.000284 |
| PBMC-2D | 0.000076 | 0.000014 | 0.000366 |

**Table 6:** Entropy gaps for PBMC-3D. Joint denotes perturbation × cell type × donor. Values are fixed-split statistics, rounded to six decimal places.

| Dataset | Perturbation | Cell type | Donor | Joint |
|---|---|---|---|---|
| PBMC-3D | 0.000327 | 0.006464 | 0.009370 | 0.000116 |

## B Implementation Details

### B.1 Data Preprocessing

For each cell, let 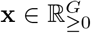 denote the raw gene-count vector and let 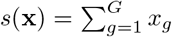 be its library size. We first perform library-size normalization to a common target sum *τ* = 10,000:

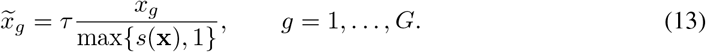

We then apply the elementwise logarithmic transform 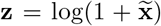. Highly variable genes are selected using only cells visible to the training split, including the corresponding training controls. For each gene, we compute its mean *µ*_*g*_ and sample variance *v*_*g*_ in this transformed space, form the log-dispersion

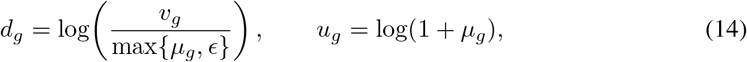

and divide genes into 20 bins according to *u*_*g*_. Within each bin *b*, we standardize *d*_*g*_ using the bin mean and standard deviation:

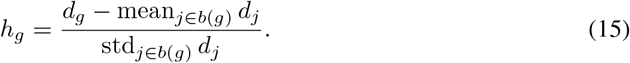

We retain the top 2,000 genes by *h*_*g*_ and use the resulting log-normalized vectors in all model inputs and losses. This train-only selection prevents test-condition expression values from affecting the feature space.

### B.2 Model and Training Configuration

Each cell is projected to a 512-dimensional representation. A Q-former with *q* = 15 learned query tokens summarizes each cell set, followed by a population-mean token. The two states in each demonstration are processed by two pair-level self-attention blocks, and target cells use four trunk cross-attention blocks to attend to the context memory. The context and target populations are sampled with *n* = 128 cells per bulk; the standard training configuration uses five context pairs for each active context group, batch size 32, and AdamW with learning rate 10^−4^ and weight decay 10^−4^. We use eight attention heads, a 2,048-dimensional feed-forward layer, zero dropout, and cosine learning-rate decay to a minimum of 10^−5^ over 40,000 steps. Each experiment takes approximately 8 hours on a single NVIDIA A100 GPU.

**Table 7:**
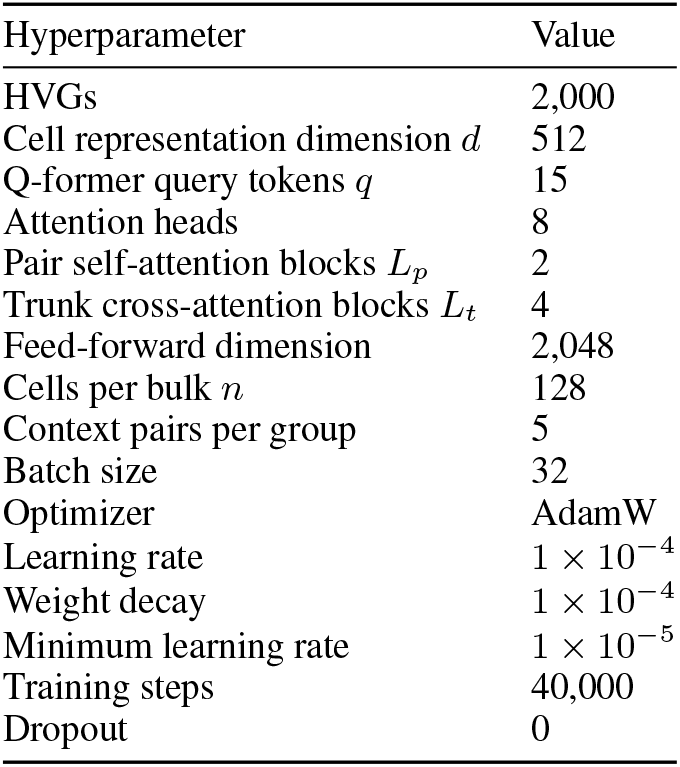
Main implementation hyperparameters.

| Hyperparameter | Value |
|---|---|
| HVGs | 2,000 |
| Cell representation dimension *d* | 512 |
| Q-former query tokens *q* | 15 |
| Attention heads | 8 |
| Pair self-attention blocks *L*<sub>*p*</sub> | 2 |
| Trunk cross-attention blocks *L*<sub>*t*</sub> | 4 |
| Feed-forward dimension | 2,048 |
| Cells per bulk *n* | 128 |
| Context pairs per group | 5 |
| Batch size | 32 |
| Optimizer | AdamW |
| Learning rate | 1 × 10<sup>−4</sup> |
| Weight decay | 1 × 10<sup>−4</sup> |
| Minimum learning rate | 1 × 10<sup>−5</sup> |
| Training steps | 40,000 |
| Dropout | 0 |

### B.2 Training Objective and Evaluation Metrics

Let 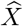, *X* ∈ℝ^*n*×*G*^ denote the predicted and observed treated cell bulks after adding the predicted residual to the target controls. We minimize the empirical energy-kernel MMD:

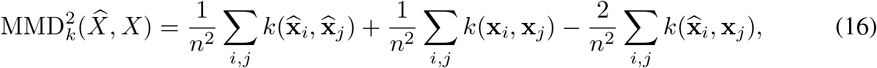

where *k*(**u, v**) = −∥**u** −**v**∥ _2_. The GeomLoss implementation uses blur parameter 0.05. Equivalently, up to the sign convention induced by this negative energy kernel and an irrelevant positive constant factor, this is the usual energy-distance combination of cross-sample and within-sample Euclidean distances. The loss is evaluated in the 2,000-gene log-normalized expression space and averaged over the batch.

For evaluation, let 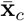 and 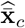 denote the observed and predicted mean expression profiles for test condition *c*, and let 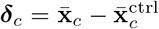 and 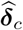 be the corresponding perturbation effects relative to the matched control mean. We report five complementary metrics, averaged over test conditions.

#### Expression error

Mean squared error and mean absolute error are

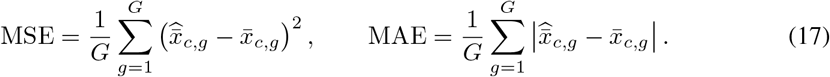

They measure absolute agreement with the observed mean profile, with lower values indicating better performance.

#### Perturbation-effect correlation

PDCorr is the Pearson correlation between the predicted and observed perturbation-effect vectors:

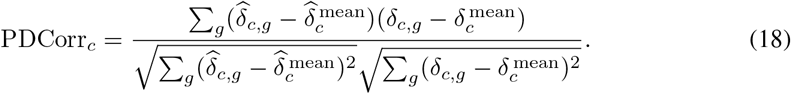

The reported PDCorr is the mean of PDCorr_*c*_ over conditions.

#### Perturbation discrimination score

For a target condition *c*, let *C*_*c*_ be the set of candidate conditions sharing all non-perturbation variables with *c*, including *c*. For each candidate *c*′ ∈ *C*_*c*_, define the cosine distance

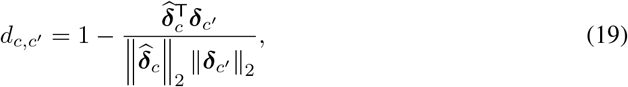

with distance 1 if either vector has norm below the numerical tolerance. Let *r*_*c*_ be the zero-based midrank of the correct candidate *c* among 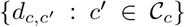, where tied candidates receive the average of their zero-based ranks. For |*C*_*c*_| *>* 1, the perturbation discrimination score is

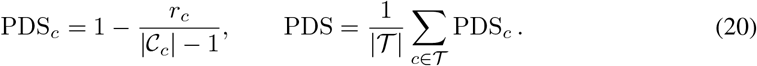

When |*C* |_*c*_ = 1, we set PDS_*c*_ = 1. Thus, higher PDS means that the prediction is closer to the correct perturbation response than to alternative responses. In computing these distances, the gene panel excludes the target gene of every candidate when that gene is present in the panel.

**Differential-expression direction reach**. Let *G*_*c*_ be the reference differentially expressed genes for condition *c*, and rank genes by the predicted statistical evidence and effect magnitude. For each prefix of this ranking, let Top_*c*_(*k*) denote the reference DE genes recovered in the prefix, where *k* = | Top_*c*_(*k*)|. Define its directional purity as

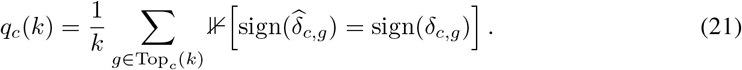

Using a fixed 90% purity threshold, the deepest qualifying prefix gives

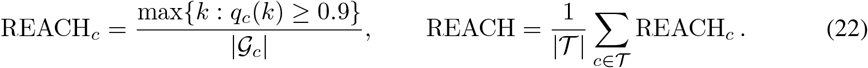

If no reference DE genes are available for a condition, its REACH value is undefined and is omitted from the macro-average; if no prefix reaches the threshold, the condition-level score is zero. REACH therefore rewards recovering a large fraction of reference DE genes with the correct response direction while prioritizing genes with stronger predicted statistical evidence. MSE and MAE are lower-is-better metrics; PDCorr, PDS, and REACH are higher-is-better metrics.

## C Additional Ablation Results

All values below are reported as mean ± standard deviation over the three seeds used for the corresponding figure. The ablations are evaluated on PBMC-3D; metrics are PDCorr, PDS, and REACH (higher is better).

### C.1 Embedding Ablation

**Table 8:**
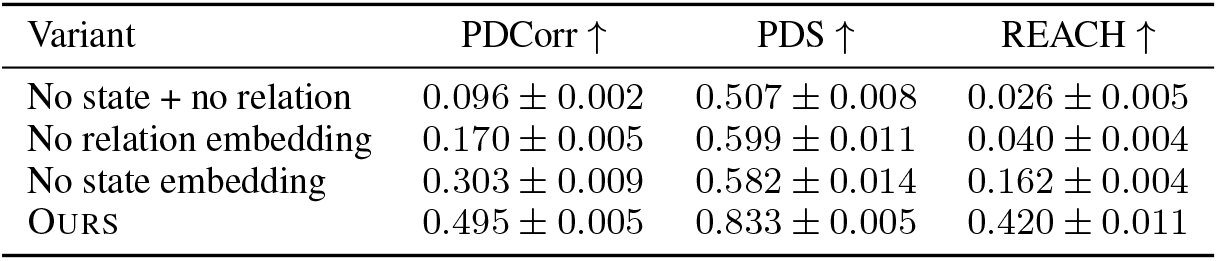
Ablation of state and relation embeddings.

| Variant | PDCorr ↑ | PDS ↑ | REACH ↑ |
|---|---|---|---|
| No state + no relation | 0.096 ± 0.002 | 0.507 ± 0.008 | 0.026 ± 0.005 |
| No relation embedding | 0.170 ± 0.005 | 0.599 ± 0.011 | 0.040 ± 0.004 |
| No state embedding | 0.303 ± 0.009 | 0.582 ± 0.014 | 0.162 ± 0.004 |
| Ours | 0.495 ± 0.005 | 0.833 ± 0.005 | 0.420 ± 0.011 |

### C.2 Context Composition Ablation

**Table 9:**
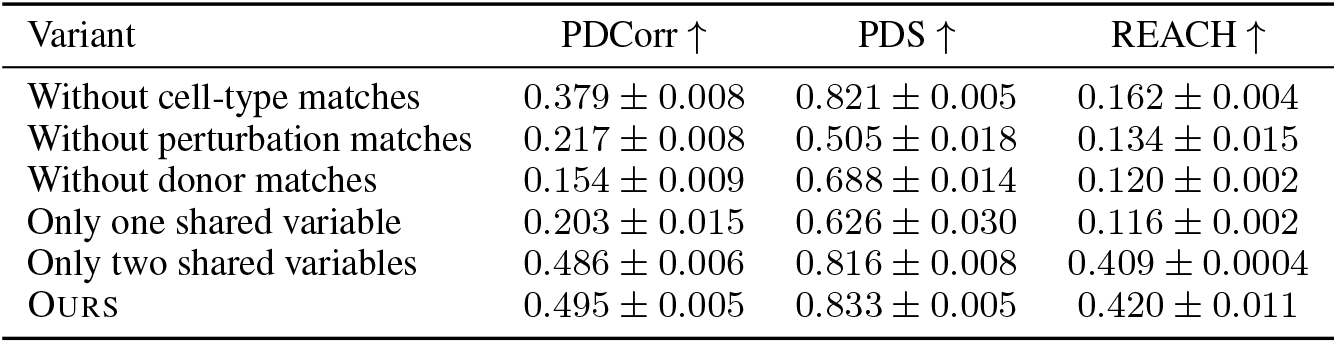
Ablation of context groups defined by variables shared with the target.

| Variant | PDCorr ↑ | PDS ↑ | REACH ↑ |
|---|---|---|---|
| Without cell-type matches | 0.379 ± 0.008 | 0.821 ± 0.005 | 0.162 ± 0.004 |
| Without perturbation matches | 0.217 ± 0.008 | 0.505 ± 0.018 | 0.134 ± 0.015 |
| Without donor matches | 0.154 ± 0.009 | 0.688 ± 0.014 | 0.120 ± 0.002 |
| Only one shared variable | 0.203 ± 0.015 | 0.626 ± 0.030 | 0.116 ± 0.002 |
| Only two shared variables | 0.486 ± 0.006 | 0.816 ± 0.008 | 0.409 ± 0.0004 |
| OURS | 0.495 ± 0.005 | 0.833 ± 0.005 | 0.420 ± 0.011 |

### C.3 Context Size Ablation

**Table 10:** Context-size ablation. *m*_1_ is the number of examples per single-shared group and *m*_2_ is the number per double-shared group.

| *m*<sub>1</sub> | *m*<sub>2</sub> | PDCorr ↑ | PDS ↑ | REACH ↑ |
|---|---|---|---|---|
| 1 | 1 | 0.423 ± 0.025 | 0.774 ± 0.014 | 0.320 ± 0.002 |
| 1 | 2 | 0.474 ± 0.013 | 0.808 ± 0.013 | 0.369 ± 0.013 |
| 1 | 5 | 0.498 ± 0.005 | 0.832 ± 0.008 | 0.420 ± 0.018 |
| 1 | 8 | 0.508 ± 0.003 | 0.847 ± 0.001 | 0.438 ± 0.004 |
| 1 | 9 | 0.509 ± 0.002 | 0.842 ± 0.005 | 0.443 ± 0.011 |
| 2 | 1 | 0.424 ± 0.021 | 0.775 ± 0.018 | 0.319 ± 0.008 |
| 2 | 2 | 0.476 ± 0.012 | 0.814 ± 0.011 | 0.378 ± 0.007 |
| 2 | 5 | 0.497 ± 0.006 | 0.834 ± 0.007 | 0.415 ± 0.026 |
| 2 | 8 | 0.508 ± 0.003 | 0.846 ± 0.002 | 0.437 ± 0.004 |
| 2 | 9 | 0.509 ± 0.002 | 0.841 ± 0.004 | 0.445 ± 0.008 |
| 5 | 1 | 0.420 ± 0.020 | 0.780 ± 0.017 | 0.313 ± 0.014 |
| 5 | 2 | 0.472 ± 0.013 | 0.814 ± 0.014 | 0.369 ± 0.018 |
| 5 | 5 | 0.495 ± 0.005 | 0.833 ± 0.005 | 0.420 ± 0.011 |
| 5 | 8 | 0.506 ± 0.004 | 0.848 ± 0.003 | 0.436 ± 0.006 |
| 5 | 9 | 0.507 ± 0.002 | 0.843 ± 0.005 | 0.442 ± 0.009 |
| 8 | 1 | 0.416 ± 0.013 | 0.780 ± 0.011 | 0.305 ± 0.009 |
| 8 | 2 | 0.467 ± 0.013 | 0.809 ± 0.014 | 0.364 ± 0.009 |
| 8 | 5 | 0.494 ± 0.006 | 0.837 ± 0.005 | 0.417 ± 0.010 |
| 8 | 8 | 0.505 ± 0.003 | 0.847 ± 0.003 | 0.433 ± 0.004 |
| 8 | 9 | 0.506 ± 0.002 | 0.845 ± 0.003 | 0.452 ± 0.008 |
| 9 | 1 | 0.412 ± 0.020 | 0.782 ± 0.021 | 0.307 ± 0.003 |
| 9 | 2 | 0.467 ± 0.013 | 0.810 ± 0.011 | 0.363 ± 0.024 |
| 9 | 5 | 0.493 ± 0.006 | 0.836 ± 0.006 | 0.417 ± 0.017 |
| 9 | 8 | 0.504 ± 0.003 | 0.847 ± 0.003 | 0.430 ± 0.007 |
| 9 | 9 | 0.506 ± 0.002 | 0.844 ± 0.004 | 0.443 ± 0.005 |

### C.4 Random Context Replacement

Table 11 reports the numerical values underlying Figure 4b

**Table 11:** Random replacement of context pairs.

| Random context | PDCorr ↑ | PDS ↑ | REACH ↑ |
|---|---|---|---|
| 0% | 0.495 ± 0.005 | 0.833 ± 0.005 | 0.420 ± 0.011 |
| 20% | 0.422 ± 0.004 | 0.793 ± 0.003 | 0.108 ± 0.005 |
| 40% | 0.344 ± 0.004 | 0.751 ± 0.016 | 0.086 ± 0.001 |
| 60% | 0.263 ± 0.014 | 0.691 ± 0.006 | 0.069 ± 0.007 |
| 80% | 0.165 ± 0.018 | 0.602 ± 0.016 | 0.060 ± 0.008 |
| 100% | 0.071 ± 0.018 | 0.504 ± 0.021 | 0.048 ± 0.010 |

### C.5 Borrowed Context Replacement

**Table 12:** Replacing the context with the complete context of another target sharing the listed variables. The matched row is the normal context baseline.

| Context relation | PDCorr ↑ | PDS ↑ | REACH ↑ |
|---|---|---|---|
| Matched context | 0.495 ± 0.005 | 0.833 ± 0.005 | 0.420 ± 0.011 |
| Share cell type | 0.077 ± 0.011 | 0.468 ± 0.012 | 0.065 ± 0.007 |
| Share perturbation | 0.203 ± 0.002 | 0.673 ± 0.007 | 0.103 ± 0.006 |
| Share donor | 0.185 ± 0.007 | 0.480 ± 0.017 | 0.113 ± 0.018 |
| Share cell type + perturbation | 0.236 ± 0.008 | 0.668 ± 0.003 | 0.139 ± 0.010 |
| Share cell type + donor | 0.217 ± 0.002 | 0.470 ± 0.010 | 0.167 ± 0.010 |
| Share perturbation + donor | 0.424 ± 0.003 | 0.807 ± 0.005 | 0.238 ± 0.001 |

## Footnotes

1 https://github.com/Yuze-e20/RELAY

2 https://www.modelscope.cn/datasets/YuzeSun/RELAY

